# A New Hyperprior Distribution for Bayesian Regression Model with Application in Genomics

**DOI:** 10.1101/102244

**Authors:** Renato Rodrigues Silva

## Abstract

In the regression analysis, there are situations where the model have more predictor variables than observations of dependent variable, resulting in the problem known as “large p small n”. In the last fifteen years, this problem has been received a lot of attention, specially in the genome-wide context. Here we purposed the bayes H model, a bayesian regression model using mixture of two scaled inverse chi square as hyperprior distribution of variance for each regression coefficient. This model is implemented in the R package BayesH.

## Introduction

In the regression analysis, there are situations where the model have more predictor variables than observations of dependent variable, resulting in the problem known as “large p small n” [1].

To figure out this problem, there are already exists some methods developed as ridge regression [2], least absolute shrinkage and selection operator (LASSO) regression [3], bridge regression [4], smoothly clipped absolute deviation (SCAD) regression [5] and others. This class of regression models is known in the literature as regression model with penalized likelihood [6]. In the bayesian paradigma, there are also some methods purposed as stochastic search variable selection [7], and Bayesian LASSO [8].

Recently, the “large p small n” problem has been receiving more attention for scientist who works with animal or plant genetics, specifically to apply in genome-wide selection studies [9].

Genome-wide selection is a approach in quantitative genetics to predict the breeding value of the individuals from a testing population based on estimates of the molecular marker effects from training population. The training population is comprised by individuals which were genotyped and phenotyped while in the testing population the individuals are only genotyped [10], [11]. Genotyping refers to obtain the genetic makeup of individuals through some technology [12] and phenotyping is a measure of some economic importance traits as yield, height and etc [11, 13].

With advent of the high throughput genotyping plataforms, nowadays is possible to define a statistical model to identify association between molecular markers and an observed phenotype. In these models, the effects of all markers are estimated simultaneously, capturing even small effects [10, 14].

In the context of genome-wide selection, many animal and plant breeders developed some bayesian regression models to make prediction of complex traits when there are more covariables than observations of response variable. In the cornerstone publication [10], Bayes A and Bayes B models were presented. In the Bayes A, the scaled-t density were used as prior distribution of marker effects, while in the Bayes B the prior distribution were modeled using a mixture of a point of mass at zero and a scaled-t density. More recently, the use of a mixture of a point of mass at zero and a Gaussian slab were purposed. This model is known in the literature as Bayes C*π* [15, 17–19].

However, there are issues in these models which should have been taken into account. The prior distribution is always influential, therefore its choice is crucial. In this paper we proposed the fit of an Bayesian regression model with mixture of two scaled inverse chi square as hyperprior distribution of variance for each regression coefficient (bayes H model). Until our knowledge, it has never reported before.

An advantage of the model is the flexibility. Depending on values chosen for hyperparameters, is possible to obtain equivalent models to (Bayes Ridge Regression and Bayes A) or even to select variable via Gibbs Sampling in a broad sense. To illustrate to application of the Bayes H model, we analyzed some simulated and real datasets.

## Materials and Methods

### Simulated Data

The aim these simulations were compare effects of prior distribution in the prediction of complex traits in some situations such as presence or absence of strong linkage disequilibrium or oligogenic or poligenic genetic architecture. The parameter settings of four scenarios generated are presented below. The phenotype were calculated using the equation described by (2).

For each scenario the predictive performance between Bayes ridge regression and Bayes H model were compared. The table (1) displays the values of hyperparameters used in mixture of the scaled inverse chi-squared distribution. Depending on the values assigned to hyperparameters, differents model can be defined. For example, using the hyperprior A (1), the model will be equivalent to Bayes A model [10]. On the other hand, the use of hyperprior B can be considered as a variable selecion model in a broad sense.

**Table 1.** Parameter settings for four simulated scenarios

| Scenario | Population * | N. individuals | N. markers | N. QTL's | Distribution of QTL's effect |
| --- | --- | --- | --- | --- | --- |
| 1 | heterogeneous stock mice | 250 | 2500 | 50 | $\text{Gamma}(3; 0, 75)$ |
| 2 | heterogeneous stock mice | 250 | 2500 | 10 | $\text{Gamma}(3; 0, 75)$ |
| 3 | random mating | 250 | 2500 | 50 | $\text{Gamma}(3; 0, 75)$ |
| 4 | random mating | 250 | 2500 | 10 | $\text{Gamma}(3; 0, 75)$ |

\* Heterogeneous stock mice were sampled from subsampling of mice dataset , which had already analyzed by [25] and available in BGLR library of R statistical software [19]; random mating were sampled from Bernoulli distribution with allele frequency equal to 0.5.

**Table 2.** Information about hyperparameters used in each component of mixture

| Hyperprior | $\nu_1$ | $s_1$ | $\nu_2$ | $s_2$ |
| --- | --- | --- | --- | --- |
| A | 5 | 0.04 | 5 | 0.04 |
| B | 5 | 0.5 | 7 | 0.002 |

The values of hyperparameters for prior distribuition for *σ*^2^ were defined as follows: degree of freedom equal to 5 and scale parameter equal to 0.1. Figure (1) shows the influence of hyperprior distribution for *τ^2^* in the marginal prior for *β_j_*. Assuming *σ*^2^ = 1, it is observed the use of hyperprior A (mixture with same scale parameters) the marginal prior resulting for *β_j_* is a t-scaled distribution. On the contrary, when hyperprior B is used the marginal prior obtaining for *β_j_* is a mixture of t-scaled distribution with the same location parameters but different scales parameters, which results a distribution with tails heavier and sharper peaks than t-scaled.

**Fig 1.** Hyperprior distribution for *τ*^2^ and marginal prior distribution for *β_j_*

## Real Data

The real dataset is comprised by 10346 polymorphic markers scored in a mice population of 1814 individuals. The phenotype measured was body mass index (BMI) [28]. The dataset were previously by [25], which further details about about heterogeneous stock mice population can be found. It is important to mention the dataset is avalaible in R package BGLR [19].

Predictions of the complex traits in the mice dataset was done in two step. First of all, a mixed model was fitted to remove the population structure and kinship effect of dataset. In the second step, Bayes H or Bayesian Ridge Regression were fitted to make predictions considering the BLUP’s predicted from mixed model as response variable.

The inference of population was based on clustering of the loadings of two top principal components obtained from to genomic relationship matrix. The clustering was done using Gaussian mixture models implemented in the library Mclust of R statistical software [23], [29] e [30]. Several mixture models were fitted and bayesian information criterion was used to select the best model. The candidate models differ each other in relation to covariance matrix of each component of mixture. The general structure of covariance matrix is Σ_*k*_ = λ*D_k_A_k_D´_k_* where Σ_*k*_ is the covariance matrix of *kth* component of mixture model, *D_k_* is the orthogonal matrix of eigenvectors, *A_k_* is the diagonal matrix whose elements are proportional to the eigenvalues and λ is a scale parameter [30].

## Phenotypic Analysis-Mixed Model

Before to predict the molecular breeding value of BMI the phenotypic analysis was done. Phenotypic analysis consisted fitting the mixed model to heterogeneous stock mice population and predict the best linear unbiased predictor (BLUP) for each individual [10,14].

The mixed model is defined by

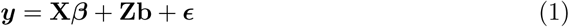

where ***y*** is the vector of response variable; **X** is the incidence matrix of the fixed effects; *β* is the vector of fixed effects that represents (litter, gender, year and population structure); **Z** the incidence matrix of random effects; **b** the vector of random effects that follows Gaussian distribution with mean 0 and variance 
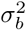
 and *ϵ* the random error.

Population structure was infered using the results from fitting of Gaussian Mixture Models implemented in the package mclust, a library of **R** statistical software [23,29,30].

## Genomic Selection-Statistical Model

The statistical model is defined by

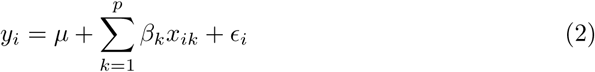

where *y_i_* is the *i* – *th* observation of response variable; *β_k_* is the *k* – *th* regression coefficient of model; *x_ik_* is a explanatory variable for *i* – *th* individual and *k* – *th* explanatory variable and *ϵ_i_* is the random error for *i* – *th* individual that follows *N*(0, *σ*^2^).

## Prior Distributions

Considering the fitting of the model, the prior distribution for intercept *μ*, is defined by

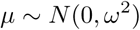

where *ω*^2^ is a hyperparameter. In practice, we used a large number for *ω*^2^ to set up this prior distribution as vague.

The prior distribution for each *β_k_* given a value of 
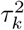
 is Gaussian, i.e,

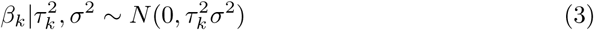

Here, is the novelty of the manuscript. The hyperprior distribution for 
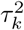
 is conditioned a latent random variable *Z_k_*. Hence, the hyperprior distribution for 
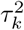
 follows the mixture of the two components scaled inverse chi square distribution, i.e

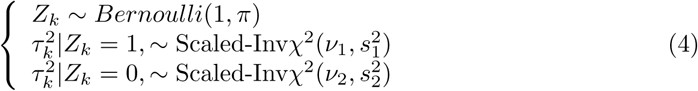

The hyperprior distribution of *π* is Beta distribution with parameters (*α*, *γ*). In pratice, we adopted *α* = 1 and *γ* = 1 to obtain a vague hyperprior.

Finally, the prior distribution for *σ* is

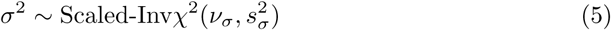

where the hyperparameters 
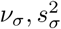
 represents the degree of freedom and scale parameters of the scaled inverse chi-square distribution.

## Likelihood and Posterior Distribution

The likelihood is defined by

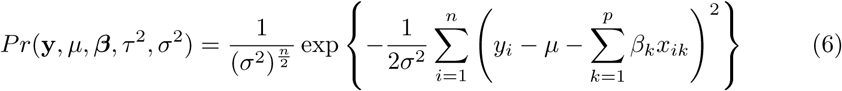

Hence, the joint posterior distribution is given by

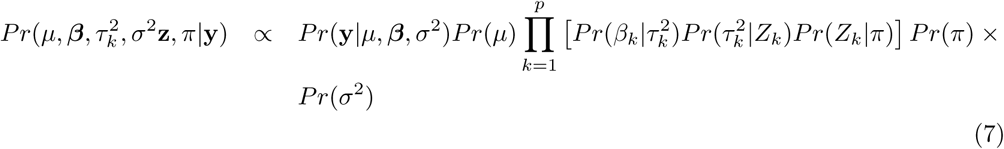

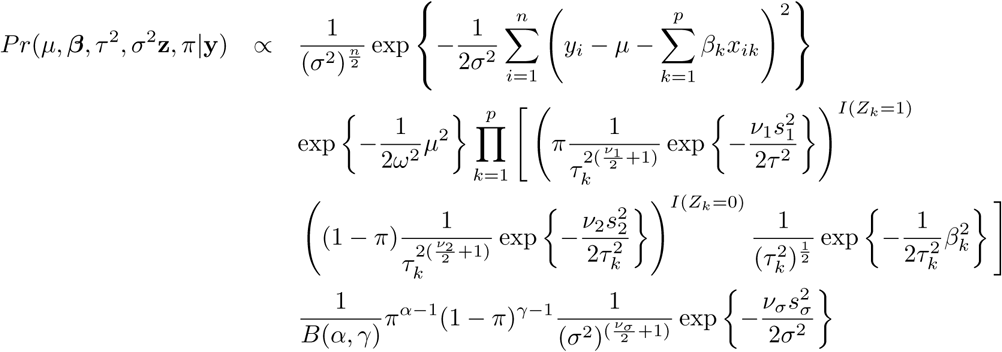

## Gibbs sampling algorithm

Gibbs sampling was used to obtain a sequence of observed values of the parameters [20], [21]. The full conditional posterior distribution for *μ* is given by

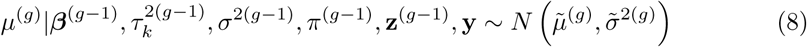

where

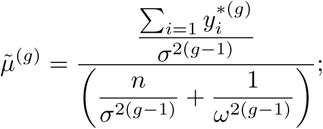

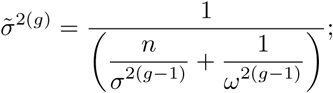

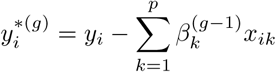

and *g* is the counter of Gibbs sampling algorithm.

The full conditional posterior distribution for 
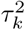
 given *Z_k_* = 1 is defined by

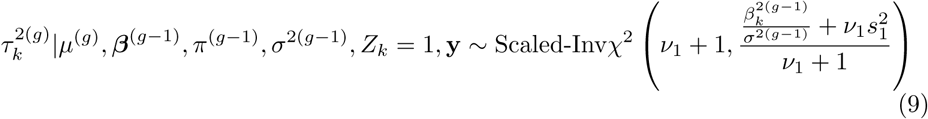

Likewise, for given *Z_k_* = 0 we have

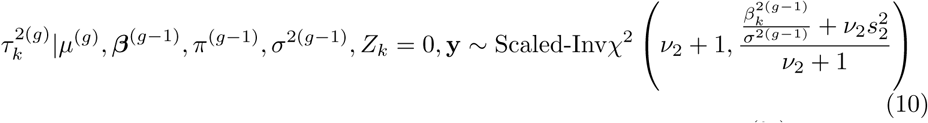

The values of *z_k_* are obtained computing the probability *Z_k_* given 
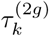
 and *π*^(*g*)^, i.e,

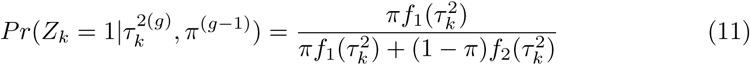

where *f*_1_(*τ*^2^) *f*_2_(*τ*^2^) are probability density functions of scaled inverse chi square distribution with parameters 
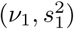
 and 
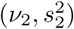
, respectively.

Moreover,

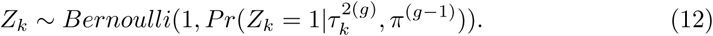

The full conditional posterior distribution for *π* is defined by

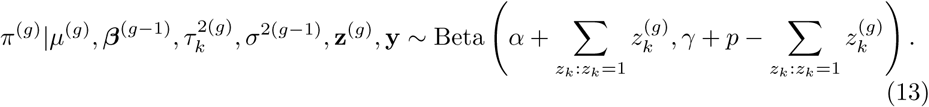

The procedure to sampling *β_k_* given *Z_k_* = 1 from the full conditional posterior distribution was adapted from the strategy purposed by [22], i.e

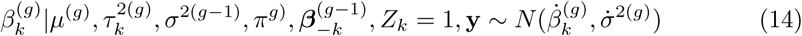

where

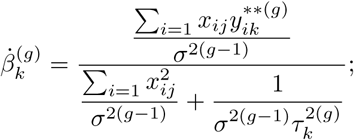

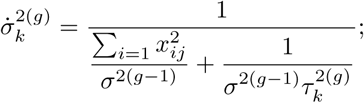

and

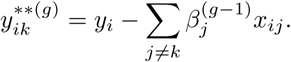

Finally, the full conditional posterior distribution for *σ*^2^ is given by

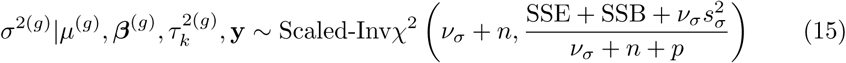

where 
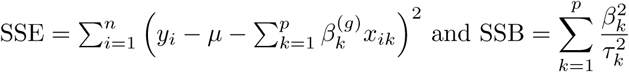
.

For Bayesian Ridge Regression, there are a unique *τ*^2^ hyperparameter. The prior distribution for *τ*^2^ and *σ*^2^ follows scaled inverse chi square with hyperparameters (*v*, *s*^2^) and 
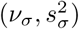
.

The full conditional posterior distribution for *τ*^2^ is

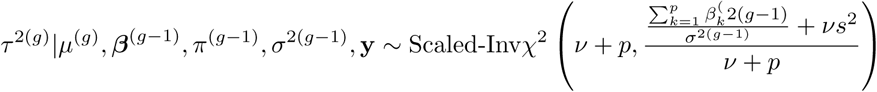

and for *σ*^2^, we have

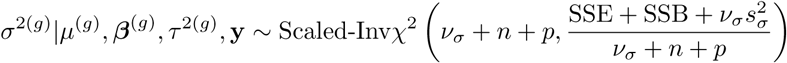

Consequently, the full conditional posterior for *β_k_* parameters is given by

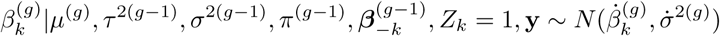

where

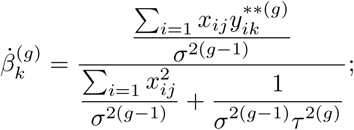

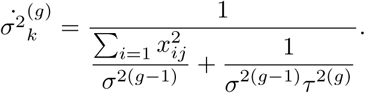

In summary, the Gibbs sampling algorithm can be defined by

For 1 to G do it:

1. Generate *μ*^(*g*)^ from (8).
2. Generate *τ*^2(*g*)^ from (9).
3. Generate **z**^(g)^ from (11) and (12).
4. Generate *π*^(*g*)^ from (13)
5. Generate each 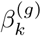 from (14).
6. Generate *σ*^2(*g*)^ from (15).

where *G* is the number of iterations.

This algorithm was implemented in a **R** package [23] called BayesH avalaible at https://cran.r-project.org/web/packages/BayesH/index.html.

## Mathematical Details about Prior Distribution

In this section we are going to show some details about conditional prior distribution for 
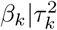
 given a prior distribuition for 
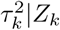
. This demonstration is based on [17]. The distribution of hyperparameter 
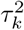
 for given *Z_k_* = 1 is described by

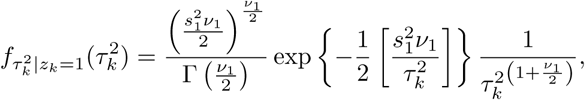

and the prior distribution for *β_k_* given *σ*^2^ and 
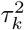
 is defined by

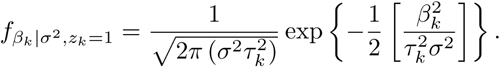

Consequently, the prior distribution for *β_k_* conditioned to *σ*^2^ and *Z_k_* = 1 is given by

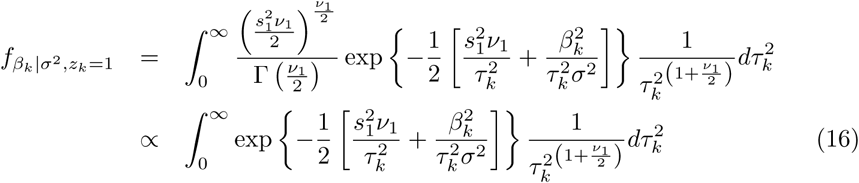

To solve the integrate written in (16) we have to make the change of variable

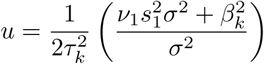

and find 
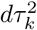 in terms of *u* and *du*. It is implies

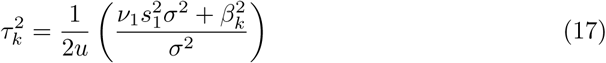

and

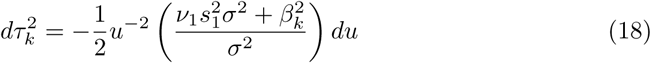

Substituing both (17) and (18) in (16) we have

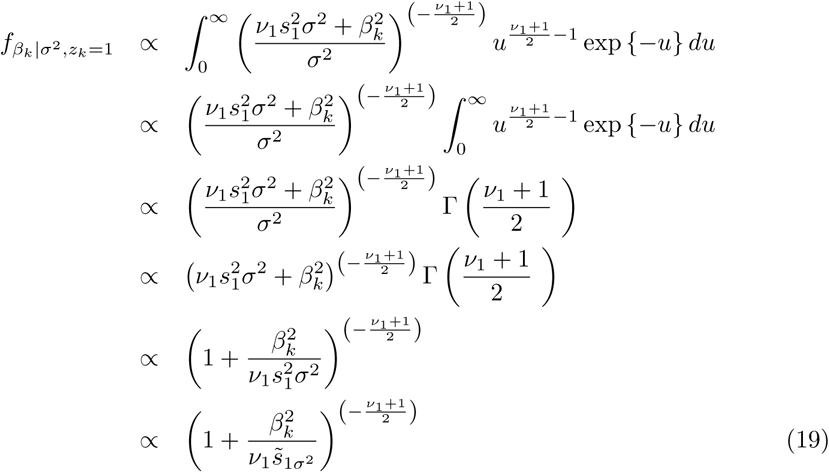

showing that (19) is a kernel of scaled t distribution [24], [17] with degree of freedom *ν*_1_ and scale parameter 
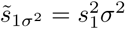
.

Likewise, for *Z*_*k*_ = 0, we have

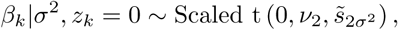

where 
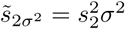
.

Consequently,

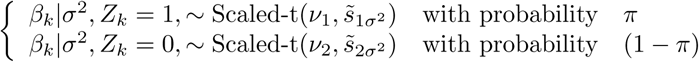

showing that for (*ν*_1_ = *ν*_2_) and 
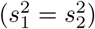
, the prior distribution for each *β_k_* of bayes H model is equivalent the prior distribuition of bayes A model. Furthermore, for 
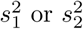
 tending to zero, the prior distribution for each *β_k_* is equivalent to bayes B model. There are other possibilities, for example, tending 
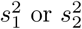
 to infinity, a mixture distribution of slab Gaussian and t-scaled distribuition is obtained as prior for each *β_k_*.

## Results and Discussion

In this study, we purposed a new hyperprior for bayesian regression model to predict complex trait. This model were applied in real and simulated datasets.

Results from cross validation studies shows the prediction accuracy of Bayes H model is slight higher than Bayesian Ridge Regression in scenarios where dataset were generated from heterogeneous stock mice population and quite higher for dataset simulated from random mating. Hence, the type population, consequently, the strength of linkage disequilibrium is more influential in the prediction than number of QTLs (2). However, comparing two datasets generated from random mating, the number of QTLs caused a increase of prediction accuracy in Bayes H model. It was not observed difference in the prediction accuracy of Bayes H when were used different hyperpriors.

**Fig 2.**
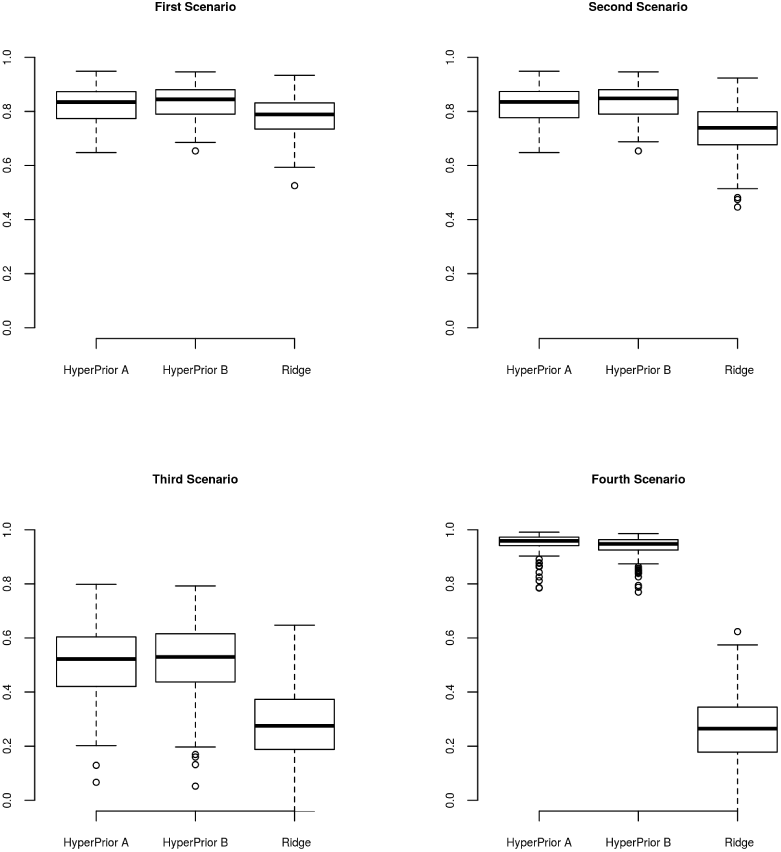
Evaluation of accuracy performance of Bayes H model using 5 fold cross validation. Box plot of Pearson’s correlation distribution between observed and predicted values for each simulated scenario.

Figure (3) shows population structure inferred from top two eigenvectors obtained from correlation matrix of mice dataset. Bayesian information criterion indicates the best model is Gaussian mixture with 8 components. Thus, we can infer the presence of eight subpopulations which are the same number of founders of heterogeneous stock mice population 3. The scatterplot of top two eigenvector estimated from to genomic relationship matrix shows the clusters 3. Prediction accuracy of BMI was compared between differents hyperpriors of Bayesian Regression models in the heterogeneous stock mice population. In order to make the comparison, 5 fold cross validation was used. Box plot of Pearson’s correlation distribution between observed and predicted values reveals moderate to high accuracy for all models 4. Moreover, the Bayesian ridge regression presented slight higher correlation in regarded to variable selecion model (hyperprior B) and quite higher correlation than model with hyperprior A.

A possible explanation of the fact that Bayes H model outperformed Bayes Ridge Regression only in a simulated dataset is the genetic architecture of the trait BMI. In the simulated data, the QTLs were the unique source of variation considered.

Furthermore, the number of QTLs used in the simulations were at most moderate (50). Therefore, models take into consideration that markers have different variances depending on their effects normally predict better than Bayesian Ridge

Regression [10,15]. Using fat percentage dataset in a dairy cattle population, which a single gene explains 50% of the genetic variation, Verbyla et. al [26] reported that predictions from fitting of Bayes Cp have more accuracy than predictions obtained by

**Fig 3.**
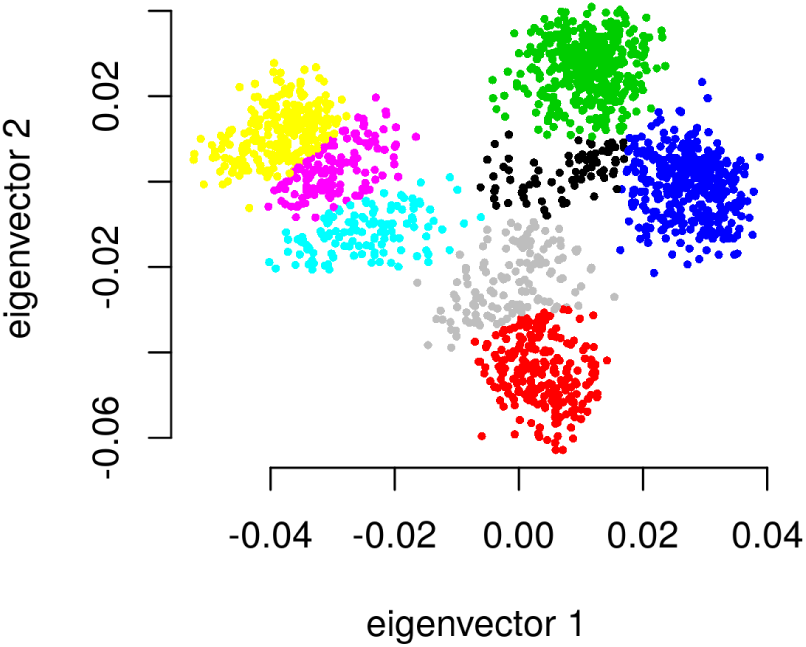
Population structure of heterogeneous stock mice.

RR BLUP. Moreover, Rezende et. al. [27] analyzed data from 17 traits measured in Pinus taeda population comprised of 951 individuals genotyped with 4853 SNPs. They concluded that for trait controlled by few genes, Fusiform rust for example, the models as Bayes A, Bayes Cp had higher predict ability in comparison to RR BLUP. On the other hand, BMI trait is considered a complex trait controlled by a large number of genes [25]. Thus, it is expected the RR BLUP would have a good predictive performance because this model considers homogeneous shrinkage of marker effects. The hypothesis is supported when we considered that genetic architecture of trait can be described by infitesimal model. However, we should have caution with these arguments, Gianola showed heuristically that Bayesian Ridge Regression or RR BLUP does not shrinkage the marker effects the same manner, the best linear unbiased predictor is sample size and allele frequency dependent [17,18]. Here we would like to speculate another hypothesis about the reason of good predictive performance of the Bayesian Ridge Regression in the real mice dataset. In the real dataset there are many source of genetic variation besides QTLS, such as: background genetic, linkage disequilibrium, epistasis effects and etc. Consequently, the linear model declared in all Bayesian model is not true. Hence, the idea to select the markers that contribute the phenotypic variation does not work well. And this case, the prediction provided by RR BLUP or Bayesian Ridge Regression would be a better approximation.

**Table 3.**
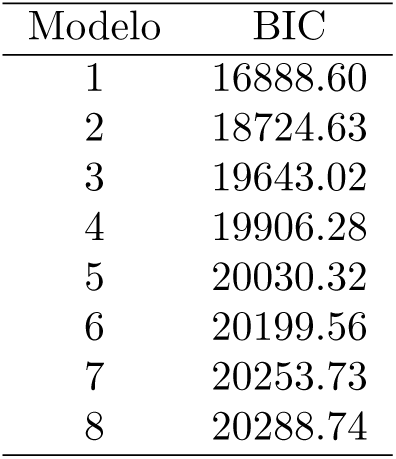
**Gaussian mixture model selection using Bayesian Information Criterion**

| Modelo | BIC |
| --- | --- |
| 1 | 16888.60 |
| 2 | 18724.63 |
| 3 | 19643.02 |
| 4 | 19906.28 |
| 5 | 20030.32 |
| 6 | 20199.56 |
| 7 | 20253.73 |
| 8 | 20288.74 |

**Fig 4.**
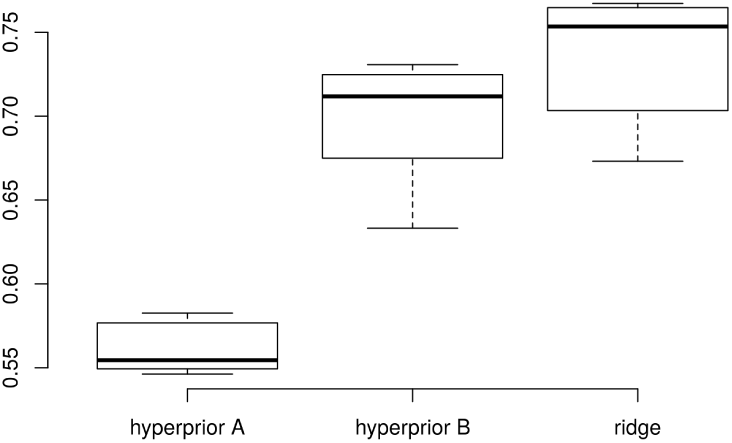
Evaluation of accuracy performance of Bayes H model using 5 fold cross validation. Box plot of Pearson’s correlation distribution between observed and predicted values for mice dataset.

